# Decoding the mystery of ultra-conservation in developmental enhancers: a role for nucleosome positioning, DNA structure and transcription factor binding

**DOI:** 10.64898/2026.07.16.739003

**Authors:** Adam Woolfe, Stephen CJ Parker, Genevieve Almouzni

## Abstract

Many human developmental enhancers are characterized by extreme evolutionary constraint in the vertebrate lineage and unique DNA sequence properties, the functional relevance of which is still unknown. Here, we investigate the consequences of their DNA sequence features on three potential aspects important for their function – transcription factor sequence recognition, chromatin accessibility and DNA structure. Using computational predictions in human as well as other vertebrates and invertebrates, we find that conserved non-coding elements (CNEs) are intrinsically nucleosome disfavoring at their core, but favor nucleosome occupancy at their borders driven by distinct nucleotide features conserved over large evolutionary distances. Nevertheless, using genome-wide nucleosome occupancy datasets, we find vertebrate CNEs exhibit higher nucleosome occupancy in comparison to surrounding regions in differentiated cells but a highly accessible conformation in embryonic tissues, suggesting a role for nucleosome positioning in their function. In addition, CNEs are exclusively enriched for homeobox transcription factor motifs, which are found at high density across their sequences. In particular, motifs specifically enriched at the boundary belong to the PBX-HOX, MEIS and POU transcription factor families, known to recognize specific DNA structural features. Consistent with this finding, CNE boundaries are enriched for unusual DNA structural motifs that may constitute a recognition mechanism by transcription factors that bind a narrow minor groove. The finding that extreme nucleotide conservation are likely to be driven by a combination of nucleosome and protein binding constraints provide a potential mechanistic insight into the function of early developmental enhancers.

## Introduction

One of the more unexpected findings following the completion of the human genome was the discovery of thousands of regions of non-coding DNA under extreme evolutionary constraint that span the entire vertebrate lineage [1, 2]. These highly conserved non-coding elements (CNEs), often several hundred base pairs in length, cluster in the genome in the vicinity of genes that coordinate early development [2]. To date, a high proportion have been shown to function as developmental enhancers that drive gene expression in a precise temporal and spatial specific manner when tested in vertebrate model embryonic systems [2, 3]. In addition, fish-mammal CNEs exhibit intriguing DNA sequence features, namely increasing G+C content towards their boundaries, followed by a sharp transition to high A+T content throughout the body of the conserved element [4].

The extreme conservation of these sequences across vertebrates extends as far as cartilaginous fish (sharks, rays and chimaeras) [5] and some as far as lamprey [6], suggesting they evolved and became fixed very early in the vertebrate lineage more than 500 million years ago. Similarly, via interspecies comparisons, invertebrates such as fly and worm have also been found to possess their own sets of CNEs [7, 8]. These CNEs, despite not sharing any sequence homology with their vertebrate counterparts, appear to be highly analogous as they also cluster around important developmental genes and share similar distinct DNA sequence properties [8]. This suggests the regulatory networks represented by CNEs are a common feature of metazoan genomes, critical in specifying a particular body plan and are likely to have evolved in parallel. Their functional importance in human health is highlighted by studies that show single point mutations and chromosomal break-points involving CNEs have been implicated in a number of congenital disorders [9]. Nevertheless, despite more than two decades since their initial discovery in the human genome, the reason for such extreme levels of conservation, their mechanism of action and the functional relevance of their DNA sequence properties remains largely unknown. Here, we focus on identifying the relevance of DNA sequence features of CNEs with regard to two aspects known to be important in the functioning of cisregulatory elements: nucleosome occupancy and transcription factor binding.

Nucleosome occupancy (the likelihood that a region of DNA will be bound by a histone octamer in a population of cells) is thought to play an important role in genome function through the control of accessibility of DNA binding proteins at promoters [10] and distal regulatory elements [11] as well as exon recognition in splicing [12]. In many cases, in vivo high or low nucleosome occupancy at a genomic locus is to some degree determined by the underlying DNA sequence [13]. It is also known that nucleosome occupancy is under the control of chromatin remodelers, in particular at sites which intrinsically disfavor nucleosome positioning [14]. The development of high throughput sequencing technology has presented the opportunity to study in-vivo nucleosome occupancy in the genome at high resolution. Here, we take advantage of the recent availability of a number of nucleosome positioning maps in human cells, as well as other vertebrate and invertebrate genomes to explore the relationship between the DNA sequence features of CNEs in metazoan genomes and their effect on intrinsic nucleosome positioning and in-vivo occupancy. In addition, using recently available high quality binding site preferences for the majority of transcription factor families in the human genome, we investigate whether CNEs represent binding sites for specific families of proteins. Finally, given the known role of DNA structure (that is, the shape of the DNA double helix, in particular the shape of the minor groove) on specific recognition by transcription factors [15], we investigate whether the distinct CNE sequence properties at their boundaries generate structural features relevant to their likely function as transcription factor binding site nodes.

## Results

### CNEs have high in-vivo nucleosome occupancy in human differentiated cells

Developmental gene targets associated with CNEs are tightly regulated and mostly inactive in terminally differentiated cell types. Recent evidence suggests that cell-type specific enhancer regions display high nucleosome occupancy in cell types in which they are not active, presumably as a lock-down mechanism to prevent their access by transcription factors [11]. We therefore began by investigating whether human CNEs, which are also inactive in differentiated cell types, also display high nucleosome occupancy. In this study, we focus on a set of over a thousand CNEs identified in comparisons between human and the Japanese pufferfish Takifugu rubripes for which sequence features have previously been characterized and whose boundaries are well defined [4].

We used genome-wide in-vivo nucleosome occupancy maps in four human differentiated cell types (CD4+, CD8+, Granulocytes and IMR90 cells), derived from mapping high coverage sequenced reads isolated from micrococcal nuclease (MNase) digested mononucleosomes (see Methods). Since the nucleosome turnover rate, nucleotide content and regulatory environment are likely to be different between intronic and intergenic regions, we also subdivided CNEs depending on whether they were intergenic (527) or intronic (499). In all cell types, we found that both intronic and intergenic CNEs had higher nucleosome occupancy inside the CNE compared to its surrounding sequence; a profile similar to that seen in a dataset of constitutively spliced exons of the same size (Figure 1A-B, Figure S1A-B), albeit at lower levels in CNEs. Studies have identified high GC content as a major contributor to high intrinsic nucleosome occupancy [16, 17]. This observation is therefore unexpected given the almost diametrically opposing mean A/T dinucleotide profiles (i.e. AA,TT, AT and TA) of CNEs and exons (Figure 1C). For intergenic CNEs, we observe higher occupancy at CNE boundaries followed by a dip in occupancy directly within the CNE in three of the four cell types we studied. Despite this, nucleosome occupancy inside the CNE is still higher than at surrounding regions. The most nucleosome disfavoring sequences are homopolymer tracts of adenines or thymines (poly(dA:dT) of at least 5bp in length, which are enriched in linker DNA and create stretches of rigidity, thereby making nucleosome wrapping energetically unfavorable [18, 19]. Surprisingly, despite their high general AT content, CNEs exhibit a significant depletion of poly(dA:dT) tracts within the body of the element compared to flanking regions (Figure 1D), similar to that seen in exons. This suggests the presence of poly(dA:dT) tracts within CNEs are generally selected against, most likely to allow the possibility of nucleosome positioning at these sites. Interestingly, the presence of poly(dA:dT) tracts are highest at the boundaries of CNEs (Figure 1D), particularly at intronic CNEs where this observation also corresponds to lower G+C content compared to intergenic CNEs (Figure S1C). Using a set of pentamer sequences previously identified significantly enriched within either nucleosomes or linkers [12], we see an increase in nucleosome-enriched pentamers just outside the CNE and an increase in linker-enriched pentamers within the CNE (Figure S1D). The changes are fairly subtle compared to exons and particularly so in intronic CNEs that show no or little change around the boundary for pentamers strongly enriched in nucleosomes or linkers.

**Figure 1:**
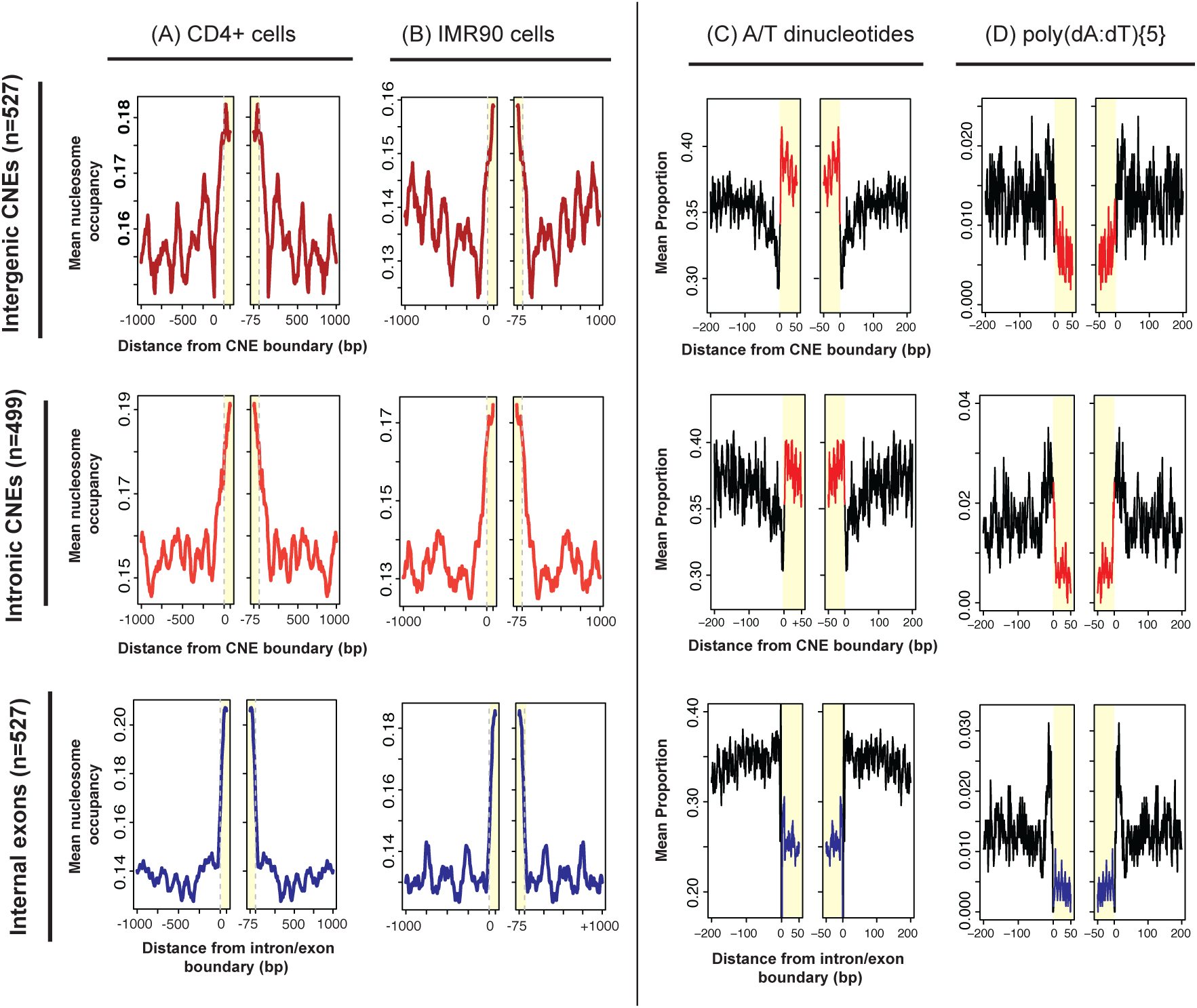
Nucleosomes are preferentially positioned at both intergenic and intronic CNEs, similar to internal exons, in different human cell types. (A-B) Mean in-vivo nucleosome occupancy in two differentiated cell types (CD4+ and IMR90) at each position 75bp internal and 1kb either side of intergenic and intronic CNEs and internal coding exons. Regions within the CNE or exon are shaded yellow. (C) Mean A/T dinucleotide composition and (D) polyA:T5 (i.e. AAAAA/TTTTT) at the same sequences.

Our results suggest that while the general sequence features of CNEs do not favor nucleosome positioning, there is evidence of selection pressure on sequence features within and surrounding CNEs that increase permissiveness to nucleosome positioning in differentiated cells in-vivo.

### CNEs have low intrinsic nucleosome positioning compared to exons and cell-type specific regulatory elements

We next investigated whether the sequence within CNEs intrinsically favor translational nucleosome positioning. Numerous studies have identified features of the underlying DNA that confer optimal interaction between histones and DNA, the most striking of which is a 10bp periodic pattern of dinucleotide frequency alternating between AT and GC dinucleotides [20]. Using computational predictions, based on these features, we confirmed high intrinsic occupancy for exons consistent with previous analyses [12], while CNEs show low intrinsic occupancy (Figure 2A).

**Figure 2:**
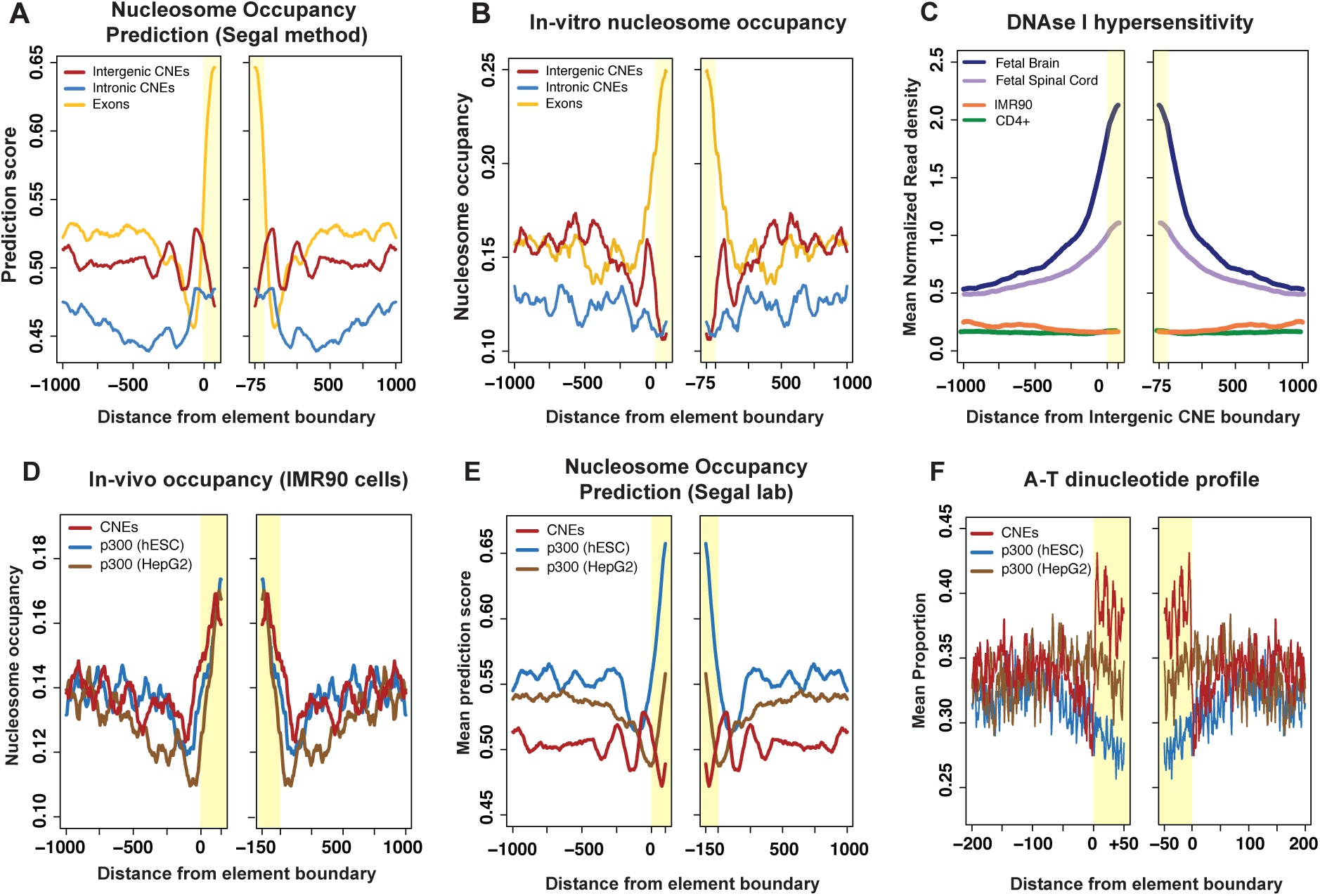
Human CNEs have low intrinsic nucleosome occupancy. (A) Nucleosome occupancy prediction based on DNA sequence alone or (B) in-vitro reconstituted nucleosomes. (C) DNAse I hypersensitivity at CNEs in human fetal tissues (brain and spinal cord) compared to two adult cells (CD4+ and IMR90). (D) In-vivo nucleosome occupancy in IMR90 cell at CNEs and inactive p300 sites from hESC and HepG2 cells. (E) Nucleosome occupancy prediction and (F) A/T dinucleotide profile of CNEs and inactive p300 sites from hESC and HepG2 cells.

We observed a difference between intronic and intergenic CNEs with intronic CNEs showing a small increase in intrinsic occupancy within the CNE, while intergenic CNEs have moderately elevated occupancy at the CNE boundaries (coinciding with elevated GC content), but low occupancy within the CNE. This difference could explain the greater level of in-vivo occupancy seen at intronic CNEs (Figure 1A-B, Figure S1A-B).

An additional approach to measure intrinsic occupancy is in-vitro reconstitution, where histones and genomic DNA are mixed and allowed to form in the absence of any chromatin remodelers and then digested with MNase. Using a previous in-vitro reconstitution dataset [21] we observed high correspondence between the pattern of nucleosome occupancy in this dataset and computational predictions for exons and intergenic CNEs (Figure 2B), confirming that CNEs, in contrast to what we observed in-vivo, have low intrinsic translational positioning. The higher predicted occupancy at intronic CNEs did not translate to higher in-vitro occupancy suggesting that the small increase in predicted occupancy is not functionally significant. In addition, we used a dataset of nucleosome-retention in human sperm, that is thought to be driven mainly by intrinsic DNA signals [16] and found very similar profiles to that seen in the in-vitro dataset (Figure S2), confirming our conclusions. These results also exclude the possibility that the in-vivo observations are due to an MNase cleavage bias that is known to prefer regions of As and Ts [22], as this bias would have been apparent in the in-vitro data too. CNEs are thought to act as developmentally important enhancers and should therefore have open chromatin conformations when they are active in development. No in-vivo nucleosome occupancy datasets are currently available at early developmental stages so we therefore used an indirect way of measuring nucleosome occupancy by studying DNase I hypersensitivity at CNEs. DNase I is an endonuclease that cleaves DNA in regions not occluded by nucleosomes representing active regions of transcription factor binding at promoters and enhancers. DNase I cleavage has been coupled with high throughput sequencing to produce genome-wide maps of active regions [23]. We measured chromatin accessibility in two human fetal tissues – brain and spinal cord (in which CNEs are known to be largely active [3]) and compared this to the same assay in two adult tissues – CD4+ and IMR90 cells – cell types for which we have already measured nucleosome occupancy. We found CNEs have significantly greater chromatin accessibility in fetal tissues than adult cells (Figure 2C) confirming that CNEs are active in development and that chromatin accessibility correlates with activity. The higher level of hypersensitivity to DNAse I that we see in fetal brain compared to fetal spinal cord may also reflect a higher number of brain enhancers compared to spinal cord enhancers in our set of CNEs. We compared in-vivo nucleosome occupancy of CNEs with cell-type specific enhancers that were inactive in our four adult cell types and found that cell-type specific enhancers have generally higher or the same nucleosome occupancy levels as CNEs (Figure 2D) depending on the cell type. This is driven by intrinsic (i.e. DNA sequence driven) nucleosome positioning through the presence of nucleosome favoring high G+C content and the absence of nucleosome disfavoring poly(dA:dT) tracts. This analysis suggests that nucleosome occupancy likely plays a similar controlling role in both cell-type specific and developmental enhancers despite significant differences in their nucleotide-content features.

### Nucleosome occupancy at CNEs across metazoan genomes

The presence of both orthologous (characterized by sequence similarity) and analogous (characterized by similar functional and sequence-based features) CNEs across metazoan genomes provides the opportunity to study whether these shared features extend to similar nucleosome occupancy profiles.

We chose metazoan genomes for which there was MNase-seq data available including three widely divergent vertebrates (human, mouse and Medaka fish) as well as fruit fly and worm. We compared in-vivo nucleosome occupancy profiles with computational predictions as well as A/T dinucleotide and poly(dA:dT)5 tract occurrences to see how sequence features affected nucleosome occupancy. We found that vertebrate and fly CNEs share similar A/T dinucleotide profiles within the body of the CNE, whereas the 200bp flanking regions show a sharp and continuous decrease in A/T dinucleotides levels in Medaka and D. melanogaster, compared to a more localized decrease starting at around 50bp flanking the CNE boundary in human or mouse (Figure 3C).

**Figure 3:**
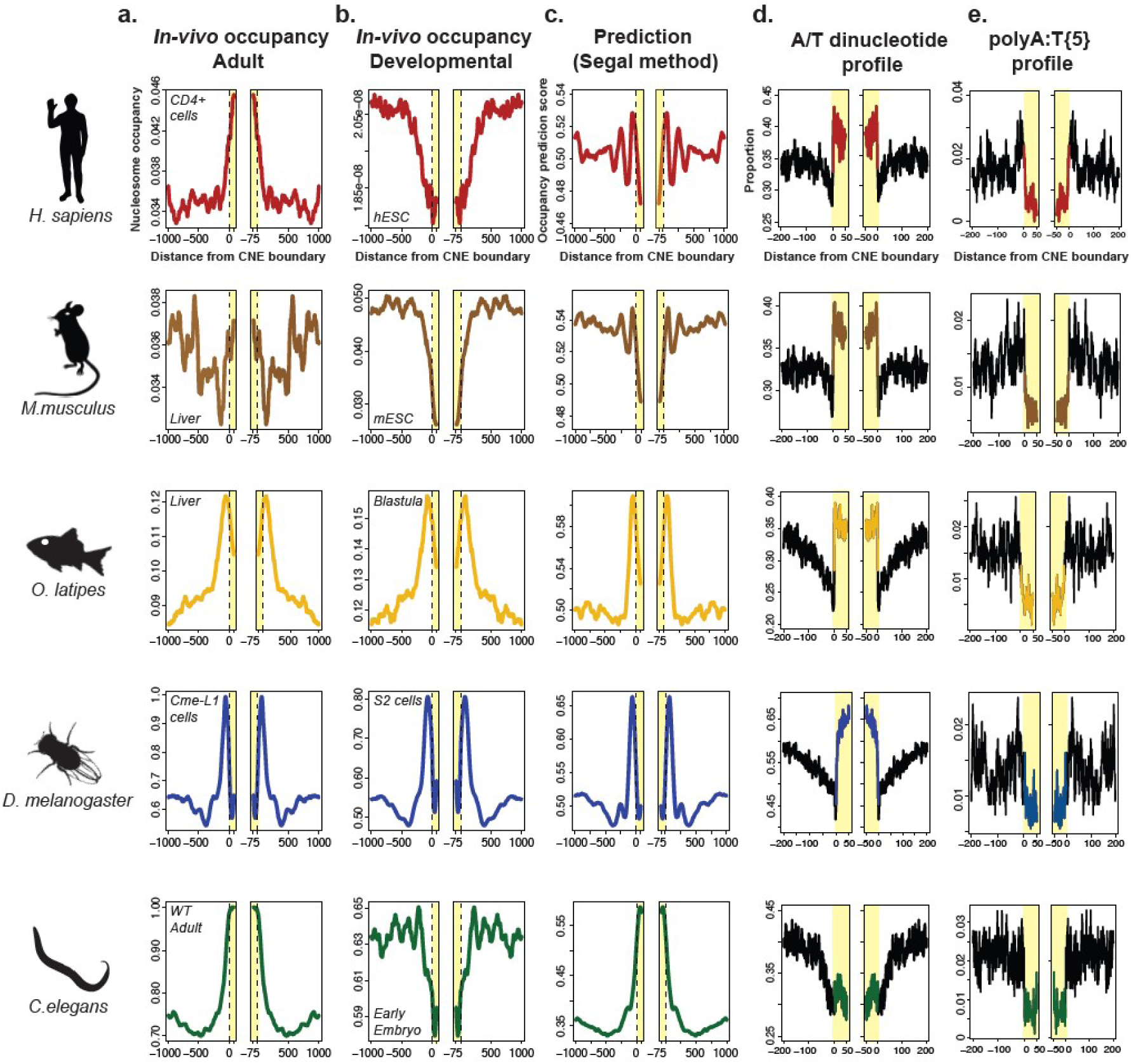
In vivo and predicted nucleosome occupancy in and around orthologous and analogous CNEs across evolution in adult and developmental cells. (a) In-vivo nucleosome occupancy in fully differentiated adult cells, (b) in-vivo nucleosome occupancy in developmental cells, (c) predicted intrinsic nucleosome occupancy using the Segal method, (d) A-T dinucleotide profile and (e) polyA:T profile at CNEs across human, mouse, Medaka fish, fruit fly and worm.

C.elegans worm CNEs (wCNEs) exhibit a profile similar to fish and flies in their flanking regions, but show only a limited increase in A/T dinucleotide content within the wCNE, and the wCNE remains fairly GC rich throughout. Nevertheless, when looking at poly(dA:dT)5 runs in CNEs, we see similar profiles throughout evolution suggesting strong evolutionary constraint against these types of sequences in distal regulatory elements in contrast to poly(dA:dT)-rich proximal promoter-based regulatory regions (Figure 3C,D). In contrast to all the other phyla we investigated, wCNEs have both high intrinsic (predicted) and adult in-vivo nucleosome occupancy, most likely due to their relatively high GC content (Figure 3A,B). A comparison of the in-vivo nucleosome occupancy in adult and embryonic worm cells (Figure S3A) confirms that wCNEs have reduced nucleosome occupancy during embryogenesis consistent with their predicted role as developmental enhancers. This finding suggests a role for nucleosome positioning in their activity. In addition, in adult worms, wCNEs as well as internal exons are enriched for the histone variant HTZ1 and depleted for H3.3 (Figure S3B+C) suggesting a divergent use of this variant in worms compared to flies and vertebrates, where neither the ortholog H2A.Z nor H3.3 are found to be enriched. Strikingly, both D. melanogaster and Medaka show strong intrinsic positioning signals for nucleosomes located on either side of the CNE, with depletion of nucleosomes within the element, which is confirmed by highly similar in-vivo data (Figure 3). In contrast, mouse and human CNEs exhibit the previously described profiles, with moderate intrinsic positioning of nucleosomes on either side of CNE and depletion within, but an opposite profile in-vivo. One reason for this difference in in-vivo occupancy profiles in these two groups could be due to the fact that data used in Medaka and Drosophila derive from embryonic tissue while those of human and mouse are from adult cells. This suggests, despite similar sequence features, that nucleosome positioning signals have diverged considerably and may play a different role in the function of conserved developmental enhancers in different species.

### CNEs are likely hotspots for binding of Homeobox transcription factors

One of the most striking features of CNEs that remains unexplained is the extensive length over which extreme conservation is observed, sometimes extending to several hundred base pairs [1, 2]. It has therefore been hypothesized that CNEs act as cis-regulatory modules that are bound by large numbers of transcription factors (TFs) [2], although little is known about the mechanism of action of these sites and the identity of the TFs that bind them. We investigated these questions computationally asking whether CNEs are enriched for specific types of TFs that could explain their unusual nucleotide composition. The availability of sets of binding site preferences for human transcription factors (TFs) has expanded significantly through the use of high throughput SELEX [22] to a degree that now allows us to tackle this question in a more comprehensive fashion. We used a set of DNA binding specificities derived from 694 position weight matrices (PWMs) representing 372 non-redundant TFs, that are divided into 24 families respectively categorized by their primary DNA binding domain (e.g. homeodomain, forkhead, T-box etc.). The number of TFs that fall into each category can be found in Supplementary Table S1.

By comparing total numbers of matches of each TF in CNEs against large sets of shuffled motifs that retain the background nucleotide composition, we identified 123 PWMs representing 92 non-redundant TFs that are significantly enriched in CNEs (detailed results for each PWM can be found in Supplementary Table S2). As a comparison we found no TF binding sites enriched in random non-repetitive genomic sequences matched for size, which confirms that CNEs are likely to represent cis-regulatory modules of clustered binding sites. Remarkably, of those TFBSs that are significantly enriched in CNEs, almost all of them (96.7%) contain a homeobox DNA binding domain, representing a highly significant difference compared to their proportion in the total set of TFs (Figure 4A-B, Figure S4).

**Figure 4:**
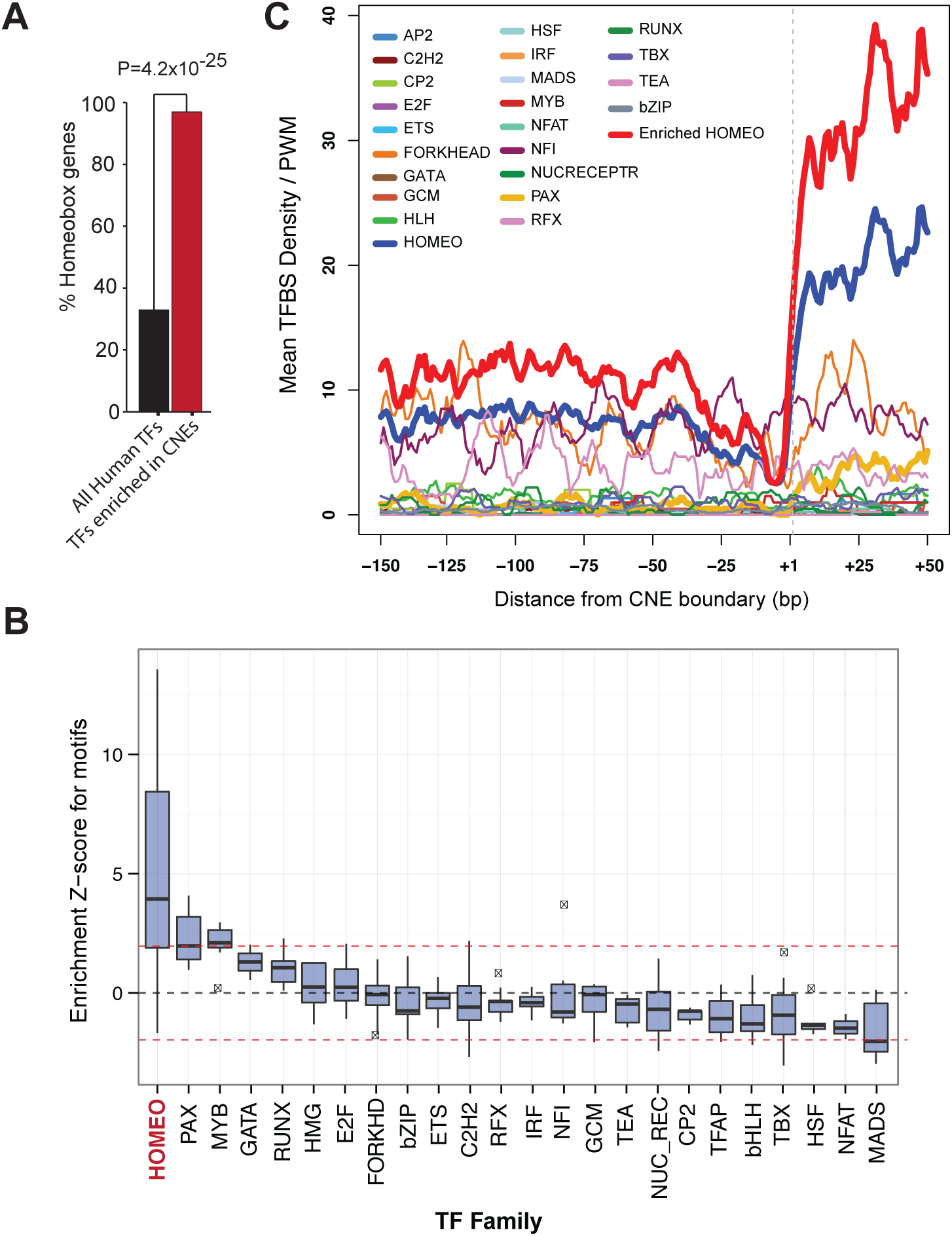
CNEs are enriched largely for and densely packed with homeobox transcription factors binding sites. (A) Histogram showing the highly significant increase in proportion of homeobox binding sites enriched in CNEs compared to the overall proportion of these matrices in the tested dataset. (B) Normalized TFBS densities for 23 families of binding site specificities for regions surrounding and within the CNEs. Densities for homeobox genes are shown as a thicker line. Densities for the subset of homeobox matrices that are significantly enriched in CNEs compared to randomized controls are show by the thick red line. A dashed grey line indicates the boundary of the CNE.

In addition, we measured the density of binding sites for each family both within and flanking CNEs, and found that CNEs contain high densities of homeobox binding sites (after normalizing for the number of PWMs per family) compared to their flanking regions (Figure 4C, dark blue line). This trend increases further when we only use those TFs that we found to be significantly enriched in CNEs (Figure 4C, red line). Homeobox TFBSs are also enriched within the flanking regions compared to the PWMs of other TF families, albeit at significantly lower densities than within the CNE, suggesting some homeobox-dependent enhancer function may be present outside the CNE core. Indeed, in contrast to exons and p300 sites, high levels of evolutionary constraint extend far from the center of the CNE (Figure S2E) providing further evidence for this hypothesis. Interestingly, homeobox binding site density decreases markedly within 40bp of the CNE boundary and is at its lowest just outside the CNE (Figure 4C) where G+C content and intrinsic nucleosome occupancy is highest. These observations suggest the presence of a boundary between the core element and surrounding regions, the relevance of which remains to be defined. In conclusion, CNEs likely constitute hotspots for AT-rich homeobox binding sites. Their presence at high density may at least partially explain the prevalence of high A+T content within the body of the element.

We used ChIP-seq studies of homeobox transcription factors in embryonic mouse cells to confirm the likelihood that these elements are indeed bound by homeobox genes. We used ChIP-seq samples for HoxA2, Meis, Pbx and Shox2 in mouse embryonic tissue to identify whether binding of these homeobox TFs to CNEs was active during embryogenesis.

All four transcription factors showed enrichment for binding moving towards the edges of the CNEs and peaking within them compared to control input samples. The degree of enrichment we observed was different across different TFs which is to be expected given that each CNE is likely to be active in very particular tissue or set of tissues that are not comprehensively covered by the tissues represented by this small sample of ChIP-seq datasets) confirming that indeed CNEs do indeed bind homeobox genes ((Figure 5A, B)). This result confirms that the enrichment of homeobox TF motif predictions are likely to represent real-life binding of these factors within CNEs.

**Figure 5:**
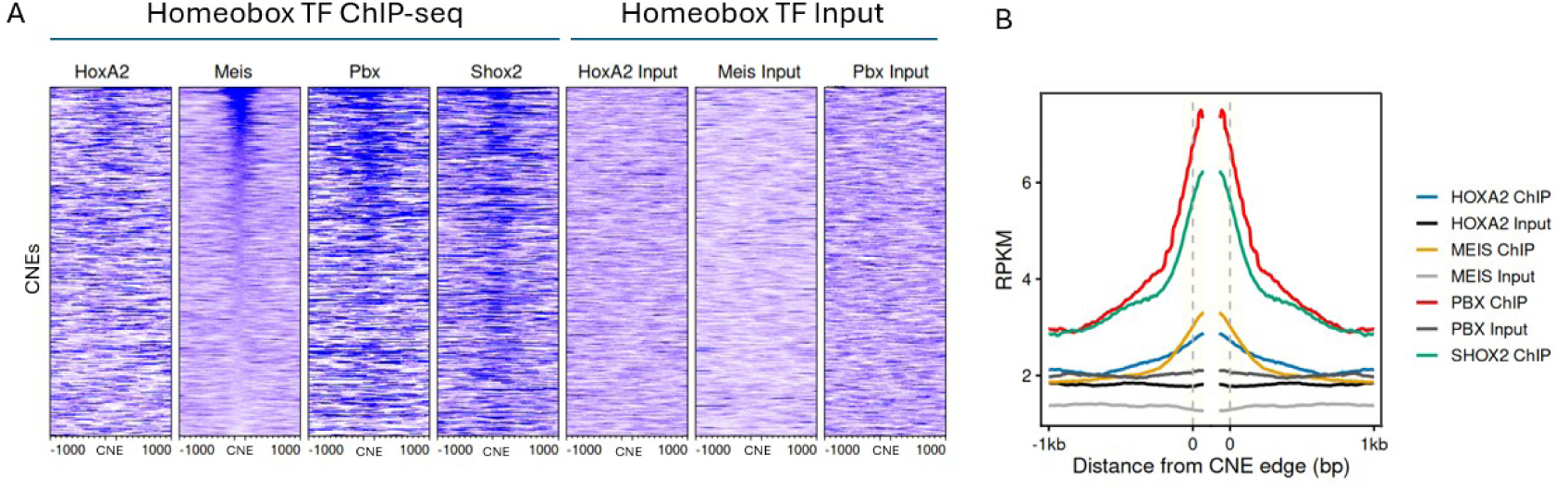
Homeobox TF CHIP-seq reveals experimental verification for Homeobox TF binding at CNEs in embryonic mouse tissue. (A) A heatmap of ChIP-seq and input read densities for four homeobox TFs (HoxA2, Meis,Pbx and Shox2 (Shox2 input was not available)) within and 1kb upstream and downstream of all CNE sequences in Mouse. (B) Averaged ChIP-seq profiles for four homeobox TFs (HoxA2, Meis, Pbx and Shox2) and matched input in mouse embryonic cells whose predicted TF matrices are enriched within CNEs (Figure S6).

Finally we used the VISTA enhancer database as the most comprehensive database of experimentally tested elements in the human genome for enhancer activity to ascertain the proportion of our set of human CNEs that are known to have enhancer activity as of April 5th 2026. We found around 31% of human CNEs have so far been experimentally tested for enhancer activity and just over 50% of those are positive for enhancer activity divided equally among intergenic and intronic CNEs (Figure S5A, B). Interestingly, the vast majority of the CNEs that were tested were active in the brain or neural tube with a smaller number in limb, eye, cranial nerve or heart. If we extrapolate from this result, assuming CNEs tested in the VISTA database are chosen at random, it suggests the likelihood that the majority of human CNEs acting as enhancers are active in neural tissue (Figure S5C). This is interesting in light of the hypothesis that CNEs represent elements setting out the fundamentals of the vertebrate body plan [24]. It appears, at least based on this incomplete experimental data, that the majority of CNEs are critical for setting out the fundamental basis of the vertebrate brain.

### CNE boundaries are enriched for binding sites of TFs that bind the minor groove

While overall A+T content is elevated in CNEs, there appears to be a particular peak in A+T content directly within the first inner 20bp, which then decreases towards the center of the element. To investigate the possible reasons for this observation we searched for TFs that were particularly enriched at the boundary of CNEs, in the vicinity of this peak and identified 3 MEIS TFs (MEIS1, MEIS2, MEIS3) that showed a particularly high density peak at the boundary than the center of the element (Figure 6A).MEIS transcription factors form complexes with HOX proteins to enhance DNA-binding specificity (e.g. [25, 26]), and TALE–Hox dimers have been shown to ‘read’ DNA shape—particularly minor-groove width—rather than just sequence [27]. Interestingly, DNA targets of another TF that also forms complexes with HOX TFs (Pbx-HOX) are also enriched in CNEs [28]. We used the PWM from this study to see whether there was a similar bias in density position and found a very similar signal (Figure 6A) suggesting that these TFs may bind similar DNA-structural features at the CNE boundary.

**Figure 6:**
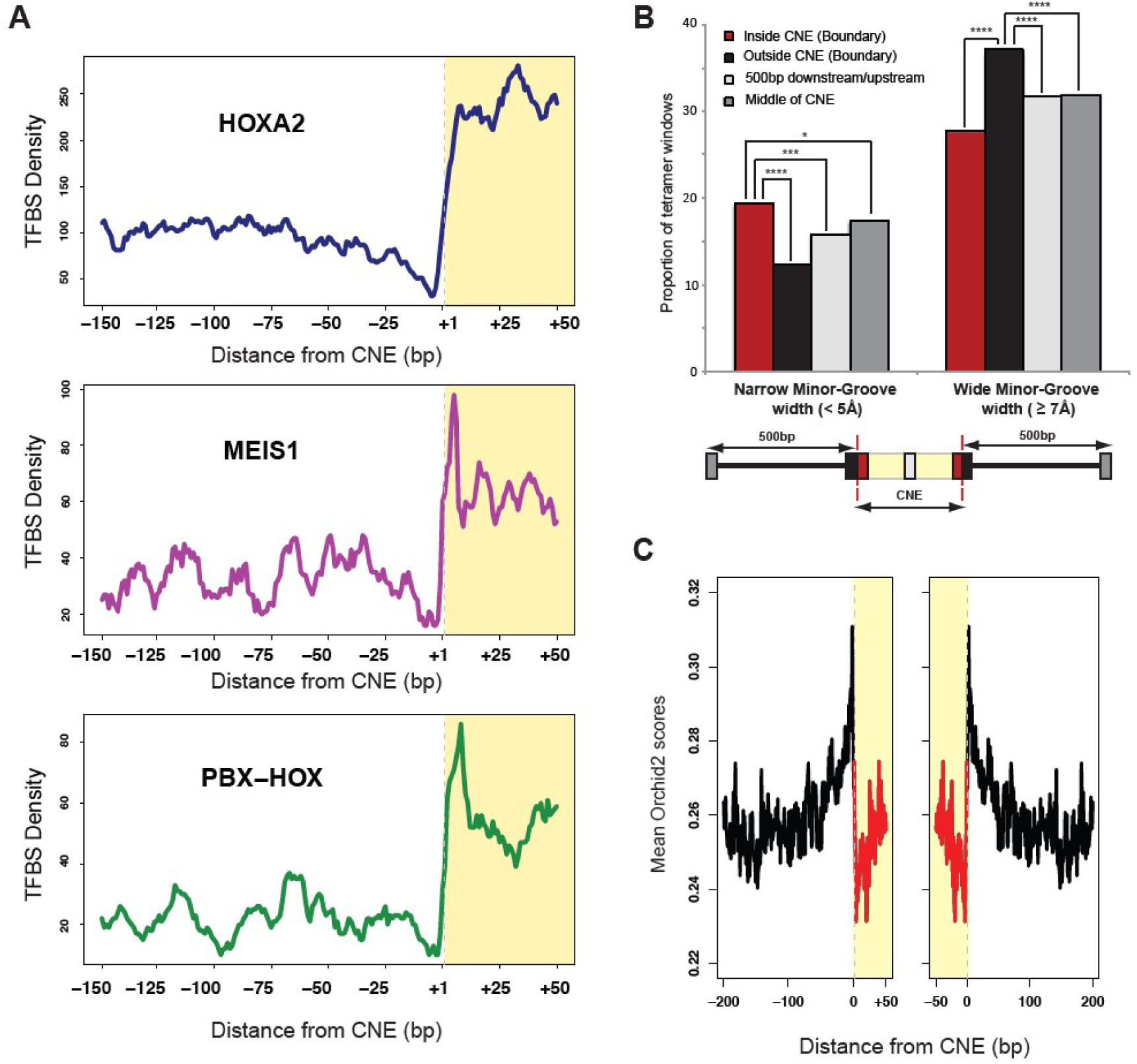
CNE boundaries are enriched for particular TF binding sites and exhibit distinct widenarrow minor-groove DNA structural motifs. (A) We identified the MEIS1 and PBX-HOX binding sites containing high densities directly at the boundary that may recognize this distinct DNA structure. A density profile of HOXA2 is shown as a comparison. (B) Sequences just inside of the boundary of CNEs are statistically enriched for 4-mers with very low minor groove widths (<5Å) while sequences just outside the boundary are enriched for 4-mers with large minor-groove widths (>7Å). Sequences in the center of CNEs and those 500bp away from the CNE boundary were used as a comparison. (C) Mean Orchid2 scores representing minor groove widths and electrostatic potential for 50bp internal and 200bp external of CNEs. Low scores represent narrow minor grooves and high electrostatic potential and high scores represent wide minor grooves and low electrostatic potential

To test whether CNE boundaries were likely to contain sequences that form regions with narrowminor groove width (MGW), we used a set of tetranucleotides known to induce narrow (<5Å) and wide (>7 Å) MGW and compared their prevalence just outside and inside of the CNE boundary as well as some distance away from the CNE and at its center. We found that there is a significant enrichment for tetranucleotides that form narrow minor grooves just inside the CNE and a significant enrichment for wide minor grooves just outside the CNE (Figure 6B).

We got similar results using the ORChID2 method for predicting hydroxyl radical (OH) cleavage intensity, which correlates with solvent accessibility of MGW (Figure 6C). OH cleavage intensity was previously shown to be predictive of MGW and electrostatic potential (Bishop et al. 2011). These observations suggest that CNE boundaries exhibit strong changes in MGW going from outside to inside the CNE that may relate to the types of TFs that bind there.

We investigated this further by studying the possible effects of the presence of these MEIS or PBX-HOX motifs at the CNE boundary.

We found that 27.4% of CNEs contain a match to either a PBX-HOX or a MEIS1 motif within the first 15bp of either edge of the CNE. Superimposition and inspection of the two peaks of motif density at the boundaries highlighted what appeared to be a shift of the PBX-HOX peak downstream of the MEIS1 peak suggesting the two motifs might localize adjacent to each other (Figure 7A). Nevertheless, inspection of individual CNEs revealed that these two motifs are almost always mutually exclusive (Figure 7B). In the 19 cases in which both motifs were found, less than 50% had both motifs on the same edge of the CNE suggesting this is not a common event. Division of the CNEs into two groups (those containing MEIS1 or PBX/HOX motifs close to the boundary, and those without) and a reanalysis of the AT dinucleotide signals, revealed that the CNEs that contained these motifs had a greater differential in GC/AT content at the CNE boundaries than those that did not contain these motifs (Figure 7D). This greater differential was also observed within the group of CNEs with the motifs, when we compared boundary regions that contained the motifs with those that did not (Figure 7E).

**Figure 7:**
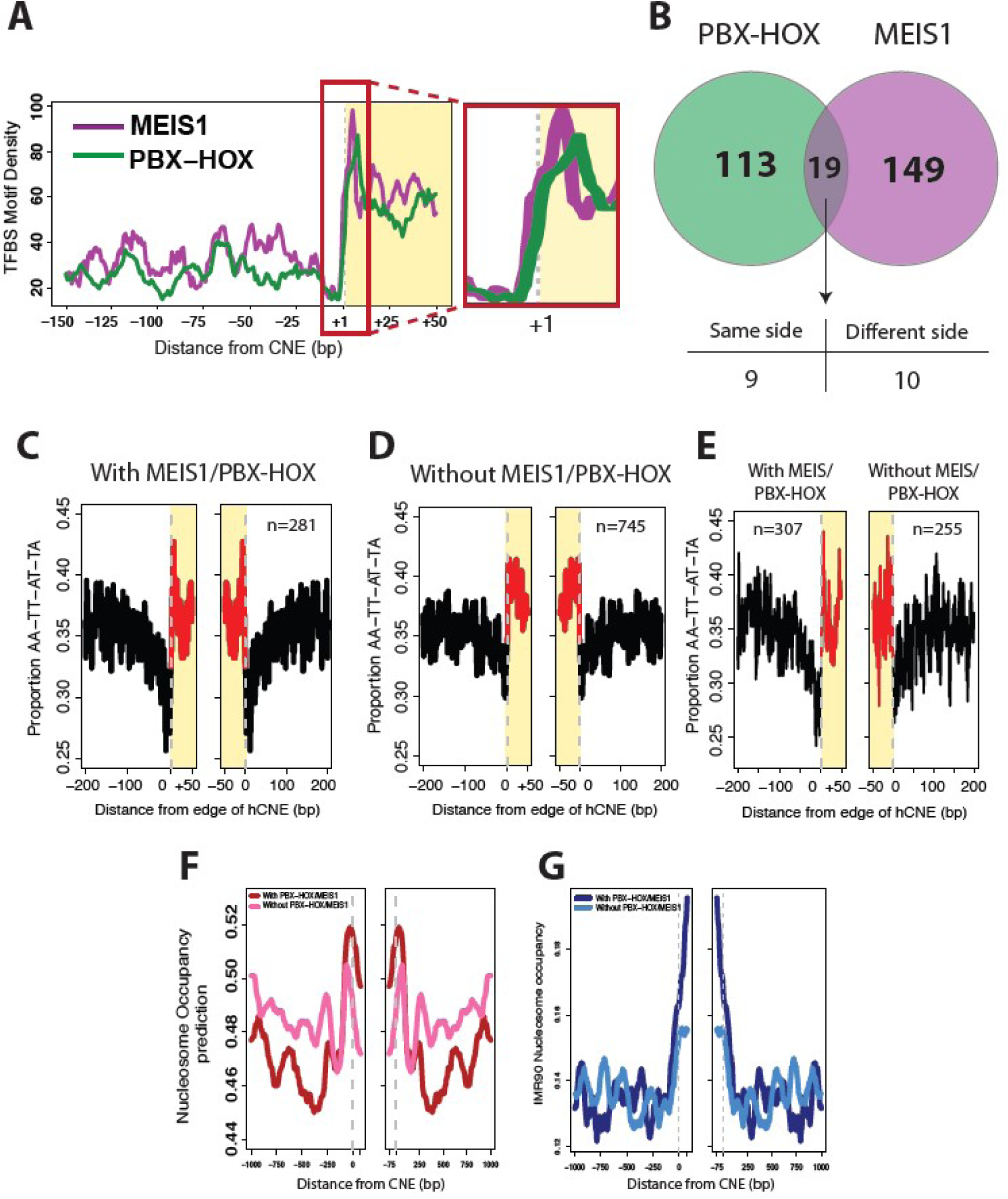
Presence of MEIS1 or PBX-HOX motifs at CNE boundaries influence their sequence and other features. (A) Superimposed profile of motif density for MEIS1 and PBX-HOX at CNE boundaries. Closer inspection of the first 15bp inside CNEs (red box) appear to show motifs adjacent to each other. (B) Venn diagram showing the number of CNEs that have match for either PBX-HOX or MEIS1 PWM at a CNE edge and how often these are found in the same CNE. Of the 19 CNEs that contain a match for PBX-HOX and MEIS1, there are roughly equal number that are located on the same side of the CNE or on opposite sides. (C) A-T dinucleotide profiles were calculated for CNEs that do or (D) do not contain a MEIS1 or PBX-HOX in the first 15bp of the CNE (either edge). (E). A-T dinucleotide plot for 281 CNEs that contain a PBX-HOX/MEIS1 match at their boundary but carried out aligning all edges containing a match on the left, and edges without a match on the right. (F) Nucleosome occupancy prediction (Segal method) or (G) In vivo nucleosome occupancy in IMR90 cells for CNEs with or without a match to PBXHOX/MEIS1 at their boundary.

We found that this greater differential in MEIS1/PBX-HOX motif-containing CNEs appears to lead to greater predicted nucleosome occupancy at the boundary regions as well as greater in-vivo nucleosome occupancy in differentiated cells (Figures 7F and 7G) compared to CNEs that did not contain these motifs at the boundary. This suggests that the presence of these TF motifs as well as recognition by other TFs and their potential requirement for specific DNA structural features may explain the strong selection for particular sequence features at CNE boundaries across evolution.

## Discussion

The extreme conservation of non-coding elements (CNEs) associated with developmental genes has remained one of the most intriguing puzzles in genomics since their discovery over two decades ago. While these elements have been demonstrated to function as enhancers in numerous experimental systems, the mechanistic basis for their extraordinary sequence constraint has remained largely unexplained. Our results suggest that this conservation is not an enigma but a logical consequence of the unusual demands placed on these elements: CNEs must simultaneously encode a chromatin landscape compatible with developmentally-timed accessibility, a high density of binding sites for homeobox transcription factors, and DNA-structural features required for recognition by minor-groove readers at their boundaries. Together these requirements impose multiple, partially overlapping constraints on every nucleotide, leaving very little room for substitution. We discuss these constraints in turn and propose a unifying model in which boundary-bound TALE-HOX complexes nucleate combinatorial homeobox binding within an A+T-rich, intrinsically remodelable core.

### Nucleosome occupancy as a developmental access switch

A central and at first sight paradoxical observation is that CNEs are intrinsically nucleosome-disfavouring consistent with their A+T-rich cores and strong depletion of poly(dA:dT) tracts [19, 11] yet show elevated *in vivo* nucleosome occupancy in differentiated cells, while becoming highly DNase-accessible in fetal tissues where they are functional. This pattern is precisely what one would expect of a sequence-encoded "access switch": a core that, in the absence of remodellers, is not strongly committed to either accommodating or excluding a nucleosome, allowing the cell type to dictate its state. In differentiated cells where the developmental targets of CNEs are silent, nucleosomes are deposited over the element likely as a lock-down mechanism to occlude transcription factor binding, in line with the high nucleosome occupancy reported at inactive cell-type-specific enhancers [11, 29, 30]. In embryonic tissues, the same sequence is rendered accessible, presumably through the action of chromatin remodellers and pioneer-like factors that exploit the very features (low intrinsic positioning, depletion of rigid poly(dA:dT) tracts) that make CNE cores energetically receptive to nucleosome eviction. The depletion of poly(dA:dT) within CNEs is particularly informative: although these tracts are the strongest known nucleosome-disfavouring sequences in the genome, their absence within CNEs argues that selection has acted not to *prevent* nucleosome formation, but to keep the locus *remodellable* neither rigidly excluded from, nor strongly committed to, a nucleosomal state. Recent work showing that combinatorial transcription factor binding targets nucleosome arrays with defined topology [31] fits naturally with this view: CNEs are sequence-encoded platforms whose chromatin state is permissive to dynamic regulation within narrow developmental windows. The sharp compositional transitions at CNE boundaries reinforce this interpretation. The G+C peak immediately flanking the element corresponds to elevated intrinsic nucleosome positioning signals and to in-vivo occupancy in differentiated cells, suggesting that boundary nucleosomes may serve to demarcate the regulatory domain and protect it from neighbouring chromatin. Comparable boundary architectures have evolved independently in vertebrates, fish, fly and worm CNEs despite a complete lack of sequence homology, indicating that this is a fundamental design principle for developmental enhancers rather than a phylogenetic coincidence. Nevertheless, the differences we see between terminally differentiated and developmental tissue in vertebrates compared to invertebrates with regard to nucleosome occupancy suggests this mechanism has nevertheless been implemented differently across evolution.

### Homeobox transcription factors and minor-groove readout at CNE boundaries

The second major constraint we identify is the near-exclusive enrichment of homeobox binding sites within CNEs: 96.7% of significantly enriched motifs contain a homeodomain, and homeobox motifs are present at remarkable density across the body of the element. This specificity is striking and identifies CNEs as bespoke platforms for homeobox-driven regulation, consistent with their genomic association with master developmental loci and with experimental ChIP-seq evidence for HOXA2, MEIS, PBX and SHOX2 binding in mouse embryos. The density and length of these arrays naturally rationalises the unusual extent of CNE conservation: when many overlapping binding sites must be preserved across a single short stretch of DNA, neutral substitutions become vanishingly rare, and the effective constraint per nucleotide can exceed that of a typical protein-coding exon [1].

A more specific constraint operates at the boundaries themselves. We find that the immediate inner edge of CNEs is enriched for tetranucleotides with narrow minor-groove width and for high ORChID2 scores indicative of high electrostatic potential, while the outer edge is enriched for wide minor-groove tetranucleotides. This sharp width transition coincides with the most prominent peaks of MEIS1, MEIS2 and PBX-HOX motif density in our analysis. The implication is direct: CNE boundaries are not simply A+T-rich, they are *shape-rich*, and the TF families that read them are precisely those known from structural and biochemical studies to interpret minor-groove geometry rather than sequence alone. HOX and TALE proteins use insertions into a narrow minor groove as a key specificity determinant [26, 32, 33], and HOX paralog-specific recognition has been shown to depend on subtle shape readout that is largely invisible to consensus motif models [27]. Our analysis therefore suggests that the pronounced compositional asymmetry at CNE boundaries is not a passive byproduct of nucleotide content but an active, selected feature: the boundary encodes a DNA shape signature read by TALE-HOX complexes.

This view also accounts for an otherwise puzzling observation. CNEs containing a MEIS1 or PBX-HOX motif within the first 15 bp of the boundary show a sharper compositional transition and higher boundary nucleosome occupancy than those without, suggesting that these boundary TFs are coupled to the local chromatin architecture. Because minor-groove width is itself a determinant of nucleosome positioning [20], the same shape features that recruit TALE-HOX binders also help define where the boundary nucleosomes will sit—a pleasing convergence of two of the constraints we observe.

### Boundary TALE-HOX binding as a nucleator for combinatorial recruitment

The functional logic that emerges from these observations is that CNE boundaries act as docking sites for TALE-HOX complexes, while the AT-rich, remodellable core acts as a substrate for combinatorial homeobox binding once the locus is opened. This is consistent with recent in vivo work showing that HOX and TALE proteins cooperate to convert a broad, low-affinity TALE chromatin platform into high-confidence, tissue-specific enhancer activation events [**?**]. In that framework, MEIS and PBX provide an early, partially nucleosome-tolerant scaffold, and the recruitment of paralog-specific HOX proteins together with tissue-restricted cofactors then specifies which subset of available sites is activated in a given embryonic territory. The boundary motifs we identify are an excellent candidate for this scaffolding step: they sit precisely where shape and chromatin features predict that the first, nucleosome-proximal binding events should occur, and they appear to be largely mutually exclusive between MEIS1 and PBX-HOX, suggesting redundant architectural roles.

This model is increasingly supported by evidence that HOX and TALE proteins, together with related developmental homeoproteins, can engage nucleosomal DNA in a pioneer-like fashion, opening compacted chromatin and thereby licensing further TF binding [**? ?**]. Pioneer factors that use minor-groove readout including, structurally, NR5A2 and the SOX family offer an instructive parallel: minor-groove anchoring appears to be a recurring solution to the problem of sequence recognition on nucleosomal DNA. Although TALE-HOX complexes have not yet been resolved on nucleosomes, the convergence of (i) minor-groove DNA shape at the boundary, (ii) high boundary nucleosome occupancy in non-permissive cells, and (iii) cooperative HOX-TALE binding profiles at developmental enhancers in vivo [25, 34**?**] all point in the same direction. Once boundary engagement has displaced or repositioned the flanking nucleosome, the AT-rich core intrinsically nucleosome-disfavouring and densely packed with homeobox motifs becomes available for combinatorial occupancy by additional homeobox proteins, plausibly including the POU2F1/2/3 factors that we also find enriched at CNE edges. The unusual length of CNEs, often several hundred base pairs, follows naturally: the element is not a single binding site but a regulatory module whose function depends on the simultaneous presentation of many overlapping homeobox motifs.

### A multiple-constraint model for extreme conservation

Taken together, our results support a model in which CNEs are subject to at least three superimposed selective pressures: (i) maintenance of an A+T-rich, poly(dA:dT)-depleted core that is intrinsically remodellable yet permissive to nucleosome deposition in non-permissive cells; (ii) maintenance of a high density of homeobox motifs along the body of the element to support combinatorial TF binding; and (iii) maintenance of distinct minor-groove shape and motif features at the boundaries that recruit TALE-HOX scaffolds and align local nucleosomes. Because almost every nucleotide contributes to more than one of these requirements, even synonymous-equivalent substitutions are likely to compromise function, and the substitution rate observed across vertebrates becomes a quantitative reflection of the overlap between constraints rather than evidence for an exotic conservation mechanism. This multiple-constraint view is consistent with the parallel emergence of analogous CNEs in distantly related metazoans [8, 35]: when a small set of biophysical requirements (remodellable cores, dense homeobox arrays, shape-defined boundaries) must be satisfied to build a developmental enhancer, convergent solutions are expected even in the absence of sequence homology.

The same framework has direct clinical relevance. Pathogenic variants within CNEs have been documented in a growing list of congenital disorders [9, 36, 37], and our results suggest these variants may act through several non-mutually exclusive routes: disruption of an individual homeobox motif, perturbation of boundary minor-groove shape and the associated MEIS/PBX-HOX docking, or alteration of the local nucleosome positioning that switches an enhancer between its accessible and locked-down states. Disease-relevant variants need not eliminate transcription factor binding outright to be pathogenic; they may instead shift the equilibrium between the developmental and adult chromatin states of a CNE, with consequences that are likely tissue- and stage-specific.

## Conclusion

We propose that the extreme conservation of developmental CNEs reflects a multi-layered code in which sequence simultaneously specifies chromatin remodellability, dense homeobox bindingsite arrays, and minor-groove shape features that recruit TALE-HOX complexes at the boundary. Within developmentally restricted windows of accessibility, boundary-bound MEIS and PBX-HOX appear well-placed to act as nucleating scaffolds, licensing the combinatorial occupancy of the ATrich core by additional homeobox and tissue-specific transcription factors. The convergent evolution of these features across metazoa, the structural biology of HOX-TALE minor-groove readout, and recent evidence for pioneer-like activity within developmental homeoprotein complexes all support this view. Direct functional tests using base-editing and high-resolution profiling of nucleosome and TF occupancy through development [38] should now make it possible to dissect the relative contribution of each layer and to predict the functional consequences of variation within these elements.

## Methods

### Collation of deeply-conserved, regulatory and exonic datasets

Conserved non-coding elements (CNEs) between Fugu and Human were taken from [2] and coordinates updated to human genome assembly GRCh38 using the liftOver tool from the UCSC Genome Browser [39]. To ensure minimal bias from other sources of nucleosome occupancy, for intergenic CNEs we filtered out all CNEs that fell within known protein coding genes as well as within promoters (defined as 1.5kb upstream of transcription start sites) and removed any CNEs that were within 1000bp of another CNE, leaving 527 CNEs. For intronic CNEs, we removed any CNEs that were located within 1000bp of an exon, leaving 499 CNEs. Due the extreme level of conservation of these sequences across vertebrates, it was possible to carry out direct conversion of the human CNEs to the mouse (mm39) and Medaka (oryLat2) genomes using the liftOver tool. In D. melanogaster, we took ‘ultra-conserved’ elements (regions of 100% identity between D. melanogaster and D. pseudoobscura with size >=50bp) from Glazov et al [7] and wCNEs (significant matches between non-coding sequence in C. elegans vs. C. briggsae) from Vavouri et al [8] and filtered them in the same way as human CNEs, except requiring no proximity to another CNE or an exon of only 300bp (rather than 1kb),due to the smaller size of non-coding regions in these species, as well as requiring a minimum length CNE of at least 75bp. Fly UCEs and wCNEs sequences were mapped to their latest genome assemblies respectively (dm3, WS200) using the BLAST program. For worm and fly CNEs, elements were divided into intergenic and intronic and 526 and 499 elements respectively were randomly picked, so they would be comparable in size to those of vertebrate CNEs. Internal coding exons from UCSC ‘known genes’ for each species were downloaded using the UCSC table browser and filtered to retain only those larger than 75bp and less than 500bp and were not located closer than 300bp from another exon. A random subset of exons of the same size as the CNE dataset was selected to measure nucleosome occupancy. P300 ChIP-seq peaks in two different cell lines were downloaded from the Gene Expression Omnibus (GEO) website: hESC (GSM602291)[40] and HepG2 (GSM935545)[41]. All P300 peaks were filtered in a similar way to CNEs, by removing all those within introns or promoters of protein coding genes, <=500bp in length and removing those located <1000bp from another P300 sites from any of the 5 cell lines.

### Measuring nucleosome occupancy

We downloaded raw reads derived from MNase-seq experiments via the Sequence Read Archive (SRA) from the following sources: Human – activated CD4+ cells (SRX000164)[42], CD8+ cells (SRX046666-9), Granulocytes (SRX046670-9), in-vitro reconstituted nucleosomes (SRX046680-84, SRX047634)[21], IMR90 cells (SRX021427)[43], sperm cells from 4 donors (SRX007049-61)[44]; Mouse – liver cells (SRX059741)[45], Medaka – blastulae (SRX001301-5)[46], Fly S2 cells (SRX021645-7)[47] and Worm – Mixed Stage,wild-type (N2) C.elegans (SRX000425-6)[48]. ChIP-seq datasets for HTZ-1 (H2A.Z) in stage L3 wild-type C.elegans (SRX059237)[49], Mnase followed by ChIP-seq of H3K27me3, H3K4me1, H3K4me3 in activated human CD4+ cells (SRX012397)[50] were obtained similarly from SRA. Indirect nucleosome occupancy was measured by using DNAse-seq data in fetal brain primary tissue, fetal spinal cord primary tissue, CD4+ primary cells and the IMR90 cell line (SRP001371)[51]. For fetal brain and spinal cord respectively, all DNAse-seq reads from all available fetal stages were pooled into a single dataset. Reads were mapped back to their respective reference genomes (Human – hg19, Mouse – mm9, Medaka – oryLat2, Fly – dm3 and Worm – ce6) using Bowtie2 [52], allowing a maximum of 3 mismatches and reporting only unique hits to the genome.

To facilitate analysis of nucleosome positioning, all reads were shifted by 37bp in the direction of the mapped read and extended to 75bp, so that the read represents the central part of the nucleosome centered at the dyad. For MNase-seq datasets, genome-wide read coverage maps were created from Bowtie alignments using the coverage() function of the ShortRead package [53] implemented in the R statistical program. Each read-coverage map was normalized by the number of million mapped reads. Average nucleosome occupancy for regions in and around CNEs was calculated by first extracting read coverage for each base for 1000bp flanking and 75bp inside the CNE, for both sides of the CNE. Read coverage was measured across each sequence in non-overlapping sliding windows of length 5bp. As the orientation of the CNEs are not known, profiles were smoothed by taking the average of each equivalent window on either side of the CNE. Total read coverage for each window was added up across all CNEs and normalized by the total number of CNEs. Average nucleosome occupancy at exons and p300 sites was measured in the same way. Wiggle files of ChIP-chip nucleosome occupancy in worm adults (GSM464565) and embryos (GSM468574) as well as the histone variant H3.3 (80mM salt) (GSM468564)[54] were downloaded from the GEO database.

### Nucleosome occupancy prediction

We applied nucleosome occupancy prediction software from the Segal lab (v3.0) [55] with default settings to each CNE dataset by extracting 5kb of flanking sequence around each CNE and calculating the probability of occupancy for non-overlapping 5bp windows across 1kb of flanking region and 50bp internal to the CNE sequence. Average occupancy probabilities were calculated across all CNEs at each equivalent window and smoothing was carried out in the same way as for real nucleosome occupancy calculations.

### Sequence and DNA structure analysis of CNEs

Average A+T nucleotide content in the 200bp flanking and the first 50bp internal of CNEs were calculated according to Walter et al [4]. Similarly we calculated proportions of AA+TT+AT+TA dinucleotides and AAAAA+TTTTT tetramers across the same sequence. In addition, we scored the mean proportion of pentamers at each position for highly enriched and moderately enriched in nucleosomes or linkers respectively using a set of 1,024 pentamers from [12] for which log-likelihood scores were assigned depending on empirically observed tendency to be covered by a nucleosome compared to a linker region.

To study possible DNA structural aspects of CNE boundaries, we scored each position 200bp upstream and downstream and 50bp internal of CNEs using the Orchid2 algorithm [56], which measures the minor groove width (MGW) across two DNA strands based on OH radical cleavage information. To study enrichment of tetranucleotides with narrow or wide MGWs in and around CNEs we used average minor groove widths from Rohs et al [15], which are calculated from protein-DNA structures in the Protein Data Bank (PDB). We studied the presence of tetramers with ‘narrow’ and ‘wide’ MGW in 5x1bp sliding windows in 4 locations in and around CNEs: 1. immediately inside the CNE. 2. immediately outside the CNE. 3. 500bp downstream,/upstream of the CNE and 4. the center of the CNE. Tetramers are defined as ‘narrow’ (MGW of < 5 Å(angstroms)) or ‘wide’ (MGW of >= 7 Å) as in [15]. For 527 intergenic CNEs, a total of 5x2x527=5270 tetramers were studied for regions 1,2 and 3 and 5x527=2635 tetramers for region 4. For 499 intronic CNEs, 4990 windows were studied for regions 1,2 and 3 and 2495 tetramers for region 4. Significance of narrow or wide minor groove width enrichment was calculated pairwise between each location using a Pearson chi-squared test with Yates continuity correction.

### TFBS analysis

We used a set of 657 human transcription factor binding site position weight matrices (PWMs) obtained from [57]. The PWMs were clustered into 23 families (e.g. Homeobox, Fork head, etc.) according to the predominant DNA-binding domain each TF contained as defined by InterPro and requiring that a family contain at least 3 independent transcription factors. For many of the TFs, there are more than one binding site as they represent specificities using DBD or the full protein. To avoid possible bias from overrepresentation of specific factors due to uneven redundancy, we created a set of 359 non-redundant matrices by taking the DBD matrix of each independent TF primarily or the full length if this was not available. A breakdown of the number of TFs in each family can be found in Supplementary Table S1 and as a pie chart in Supplementary Figure S4. Enrichment of specific families of TFBSs in all human CNEs was carried out by first scoring the total number of matches to each PWM inside the entire set of CNEs and their reverse complement using a threshold score of 0.8. This was compared against a distribution of total matches for 100 sets of randomized CNEs representing random expectation given a similar nucleotide content background. Randomization was carried out on CNE sequences using a 2nd order Markov model, which retains dinucleotide frequency and therefore models background nucleotide content more accurately. Significance was measured by calculating the *Z*-score (and associated *P* -value) of the number of matches in real CNEs versus the distribution of 100 randomized sets using

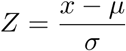

where *x* is the total number of matches in CNEs and *µ* and *σ* are the mean and standard deviation of the 100 randomized sets. A PWM was considered significant if the P-value was lower than a stringent Bonferroni corrected P-value of 0.01. Density of TFBS for each family were calculated by measuring the “coverage” of matching PWMs at each position for 200bp flanking and 50bp internal to CNEs on both sides of the CNE. Results from both sides were added together to create a single composite. Coverage was additive for all PWMs in a family of TFBSs (e.g. TBX) and then divided by the number of PWMs in that family to normalize for family size. ChIP-seq datasets for lin-39, mab-5 and egl-5 homeobox transcription factors in L3 embryonic worms were obtained from [58]. ChIP-seq of HOXA2 in mouse branchial arches at E11.5 were obtained from [34]. ChIP-seq and input of Meis and Pbx from mouse embryonic trunk (embryo lacking head, tail and limbs) was obtained from [39]. ChIP-seq of Shox2 in mouse developing limb E12.5 was obtained from [59]. ChIP-seq enriched heatmaps around the CNEs were plotted using the R library EnrichedHeatmap.

### Overlap with VISTA enhancer database

We downloaded all 4785 tested enhancer regions from the VISTA Enhancer Browser [60] as of April 6th, 2026. Coordinates for both VISTA elements and CNEs were in human genome assembly hg38. To count a CNE as intersecting a tested element we required at least 50bp of overlap. For those CNEs intersecting a positive enhancer element we extracted the tissue categories associated with that element and counted the CNE as active in that tissue if present. This means that a single element might count as being active in one or more tissues.

### Data availability

Genomic coordinates of all CNEs and exons used in this study (human, mouse, medaka, flies and worms) can be found at https://github.com/adamwoolfe/CNE_boundaries

## Acknowledgements

We thank Remo Rohs and Tianyin Zhou for help and advice on DNA structure analysis. We also thank Schraga Schwartz and Noam Field for sharing the dataset of pentamers and their scores for nucleosome favoring/disfavoring.

## Supplementary Figures

**Figure S1:**
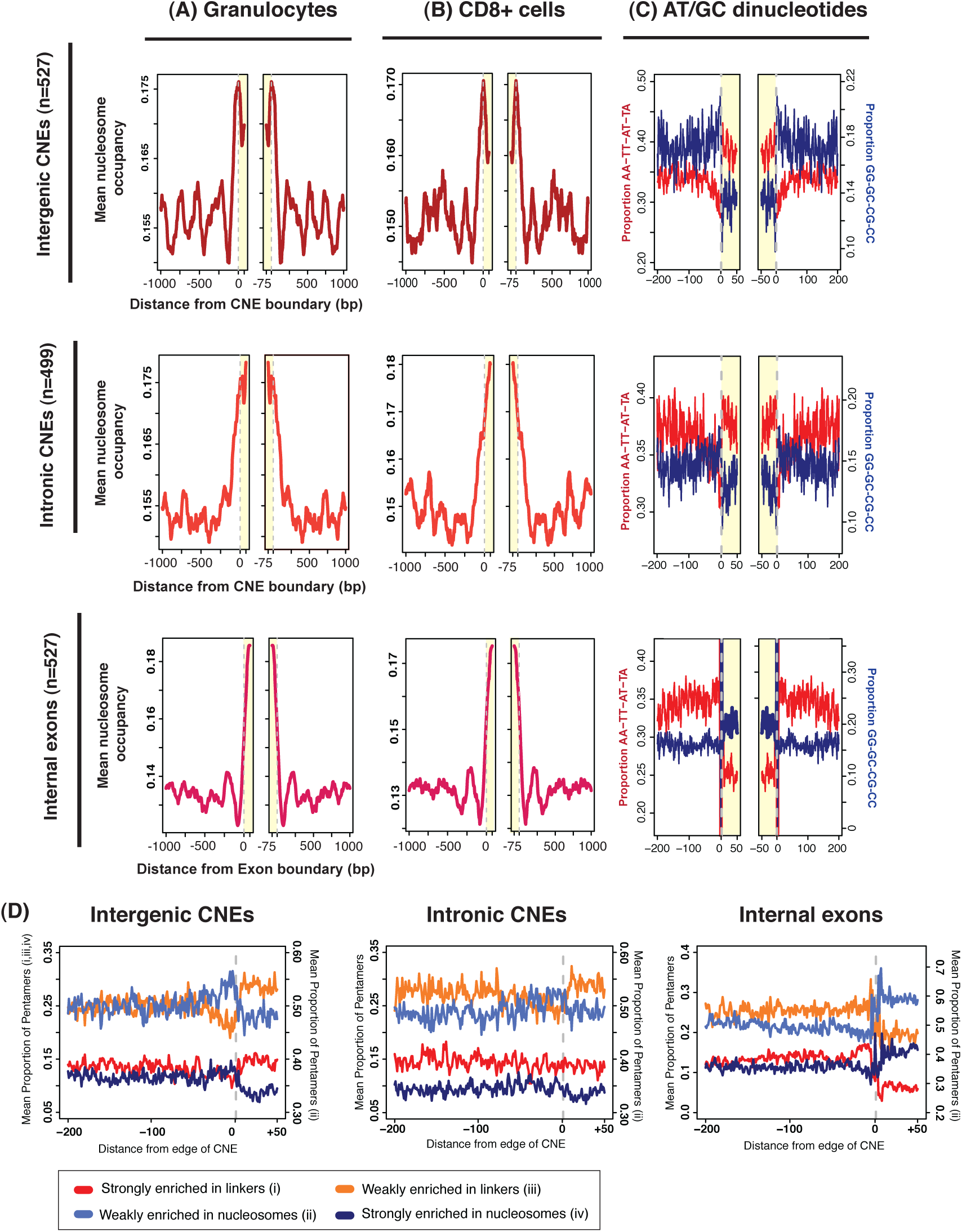
Nucleosomes are preferentially positioned at both intergenic and intronic CNEs, similar to internal exons, in different human cell types. (A-B) Mean nucleosome occupancy in two differentiated cell types (Granulocytes and CD8+) at each position 75bp internal and 1kb either side of intergenic and intronic CNEs and internal coding exons. Regions within the CNE or exon are shaded yellow. (C) Mean AT and GC dinucleotide content at the same sequences. AT dinucleotides are plotted in red and represented by values on the left hand axis while GC dinucleotides are plotted in blue and represented by values on the right hand axis. (D) Mean proportion of pentamers divided into 4 groups depending on their relative enrichment in nucleosomes or linkers genome-wide at 200bp surrounding and 50 internal to CNEs and exons.

**Figure S2:**
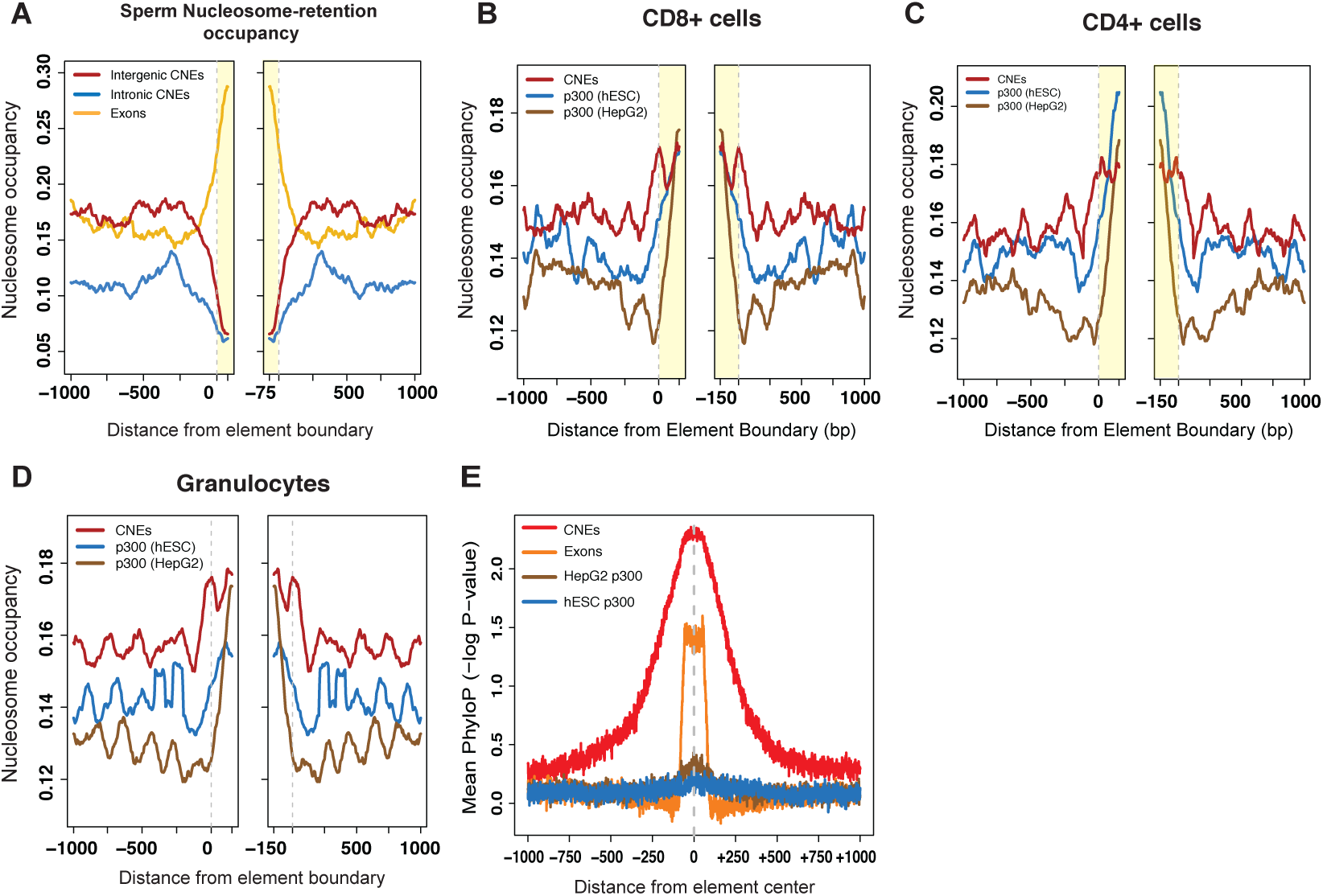
CNEs show depletion of nucleosome retention in sperm and lower occupancy than cell type specific enhancers in differentiated cells. (A) Sperm nucleosome-retention occupancy at intergenic and intronic CNEs and exons. In-vivo nucleosome occupancy in CD8+ (B), CD4+ (C) and granulocyte cells (D) at CNEs and inactive P300 sites from hESC and HepG2 cells. (E) Vertebrate-wide sequence conservation (measured using mean PhyloP score) in and around different elements showing that conservation whilst most extreme within CNEs, extends beyond their boundaries, unlike exons where conservation is limited directly to the exon itself. Small levels of conservation vertebrate-wide are seen in cell-type specific non-developmental enhancers as measured using p300 peaks in HepG2 or hES cells.

**Figure S3:**
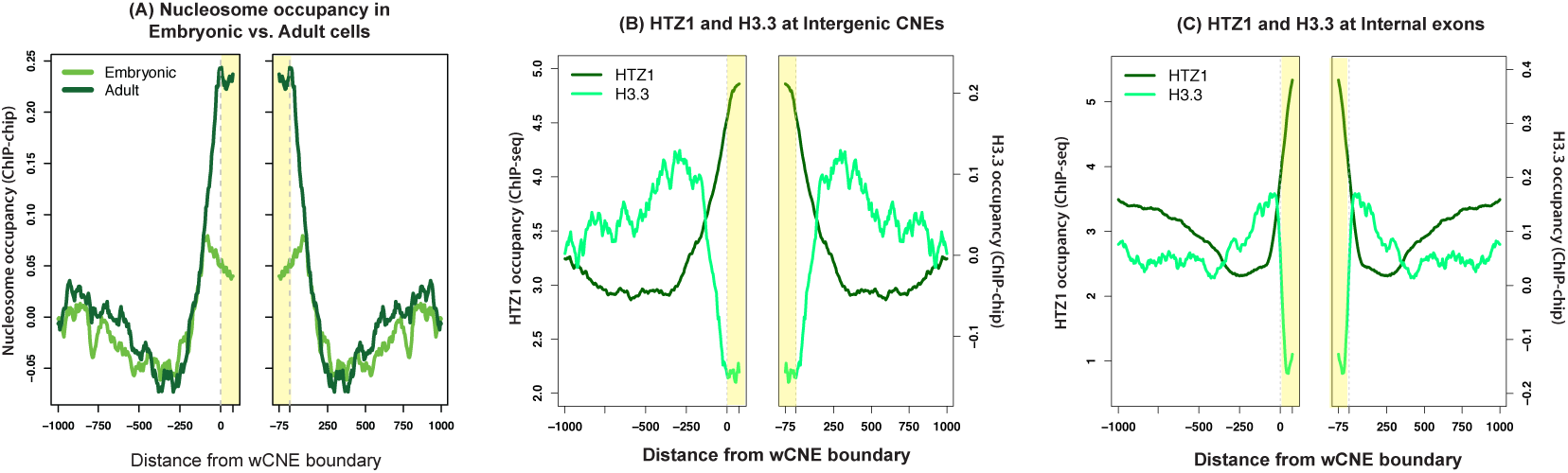
CNEs in worms have a more open chromatin conformation during development, and utilize the histone variant HTZ1 over H3.3, similar to coding exons. (A) Mean nucleosome occupancy at worm CNEs (wCNEs) in adult and embryonic cells, histone variants HTZ1 and H3.3 (B) and the same variants at exons (C).

**Figure S4:**
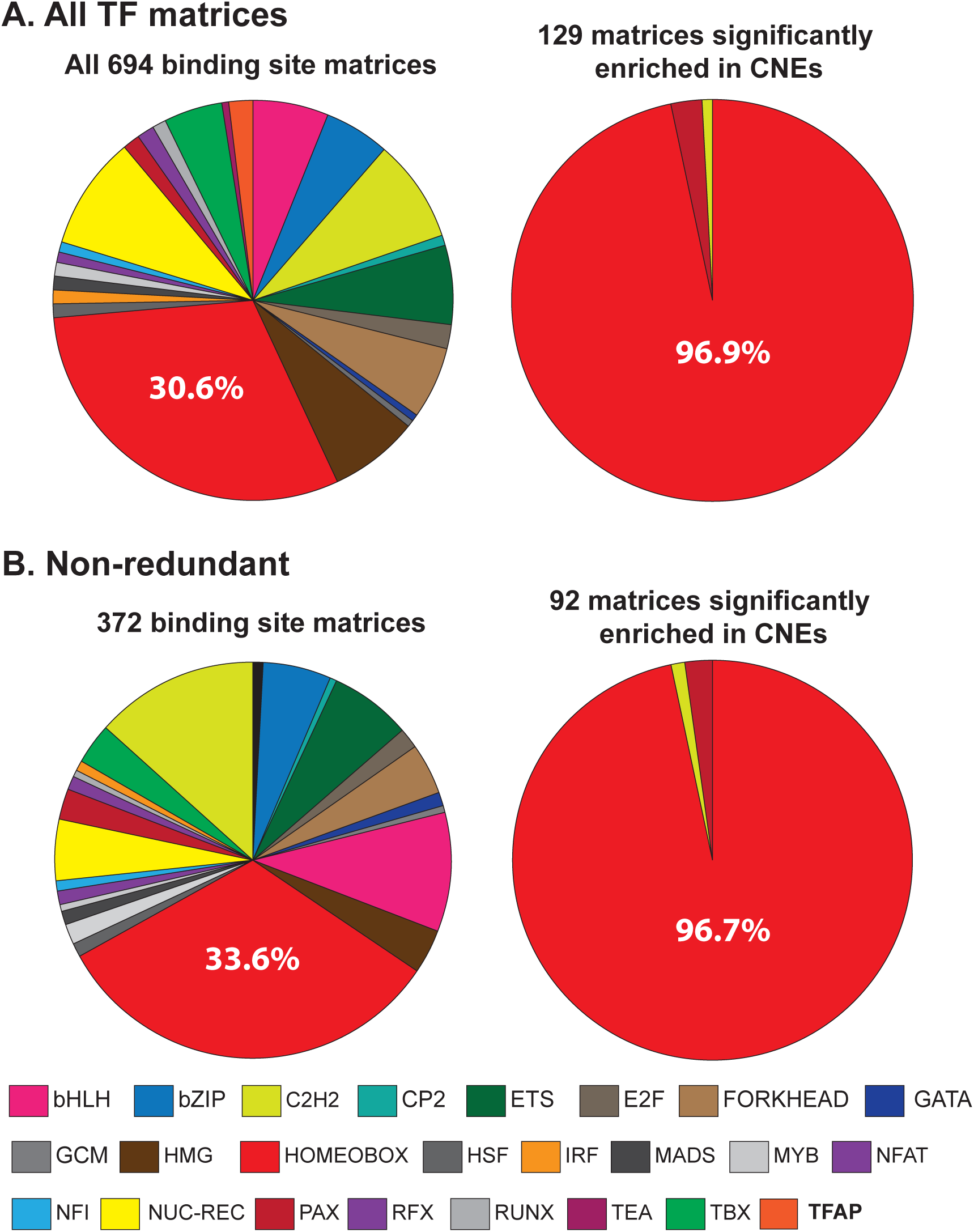
CNEs are enriched almost exclusively for homeobox transcription factor binding sites. Proportion of TF binding site matrices made up from each of 24 different types of TF families and the proportion that are significantly enriched within CNEs using the entire set of redundant matrices (A) and non-redundant matrices (B). The percentage of homeobox matrices (bright red pie section) is indicated.

**Figure S5:**
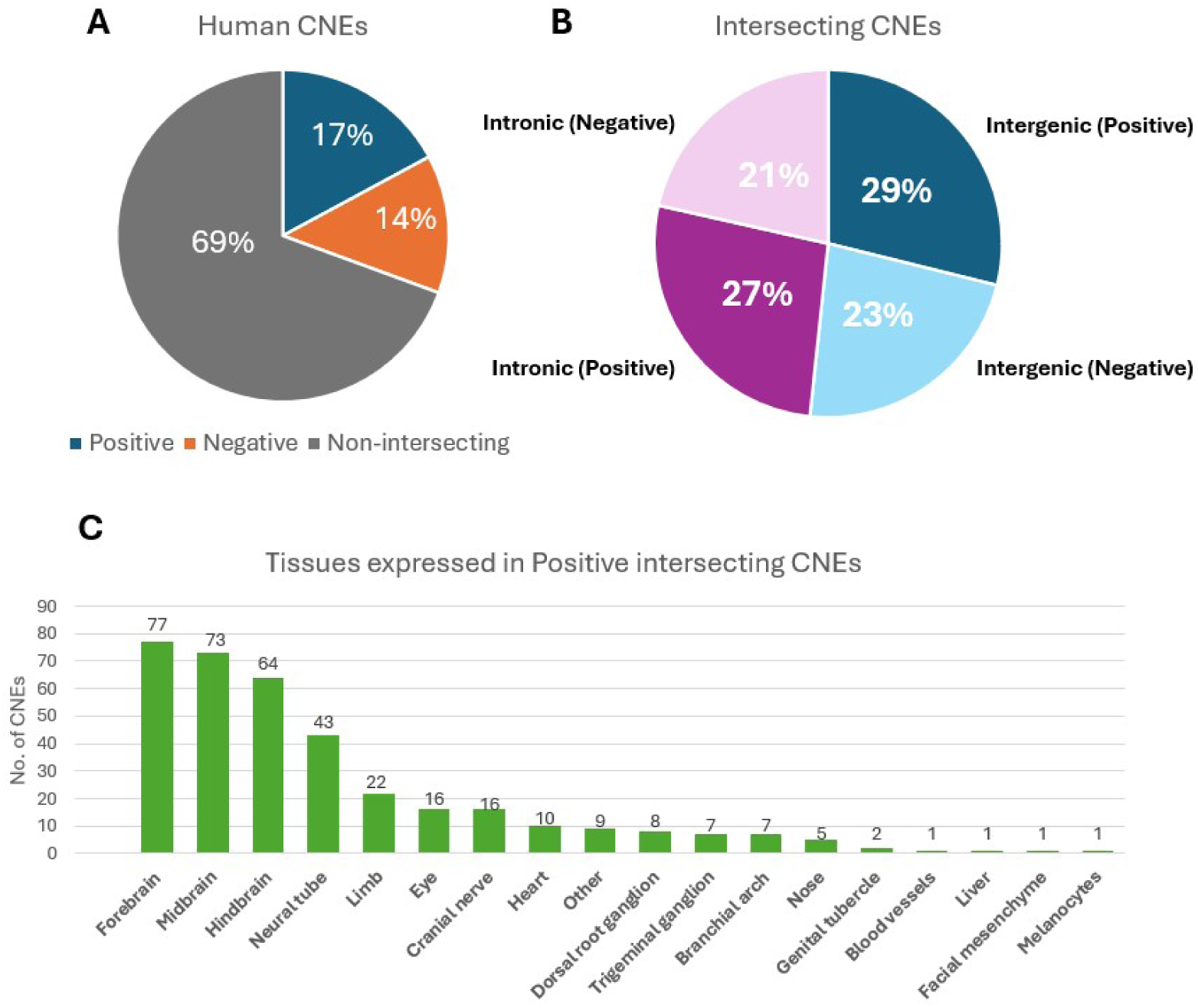
Intersection between human CNEs and experimentally tested enhancers from the VISTA database. (A) Proportion of human CNEs intersecting VISTA elements tested positive for enhancer activity, tested negative for enhancer activity or do not intersect any tested element in VISTA. (B) Proportion of positive or negative elements that fall within intergenic or intronic regions. (C) Number of intersecting human CNEs with positive VISTA elements and the organ in which they are active.

**Figure S6:**
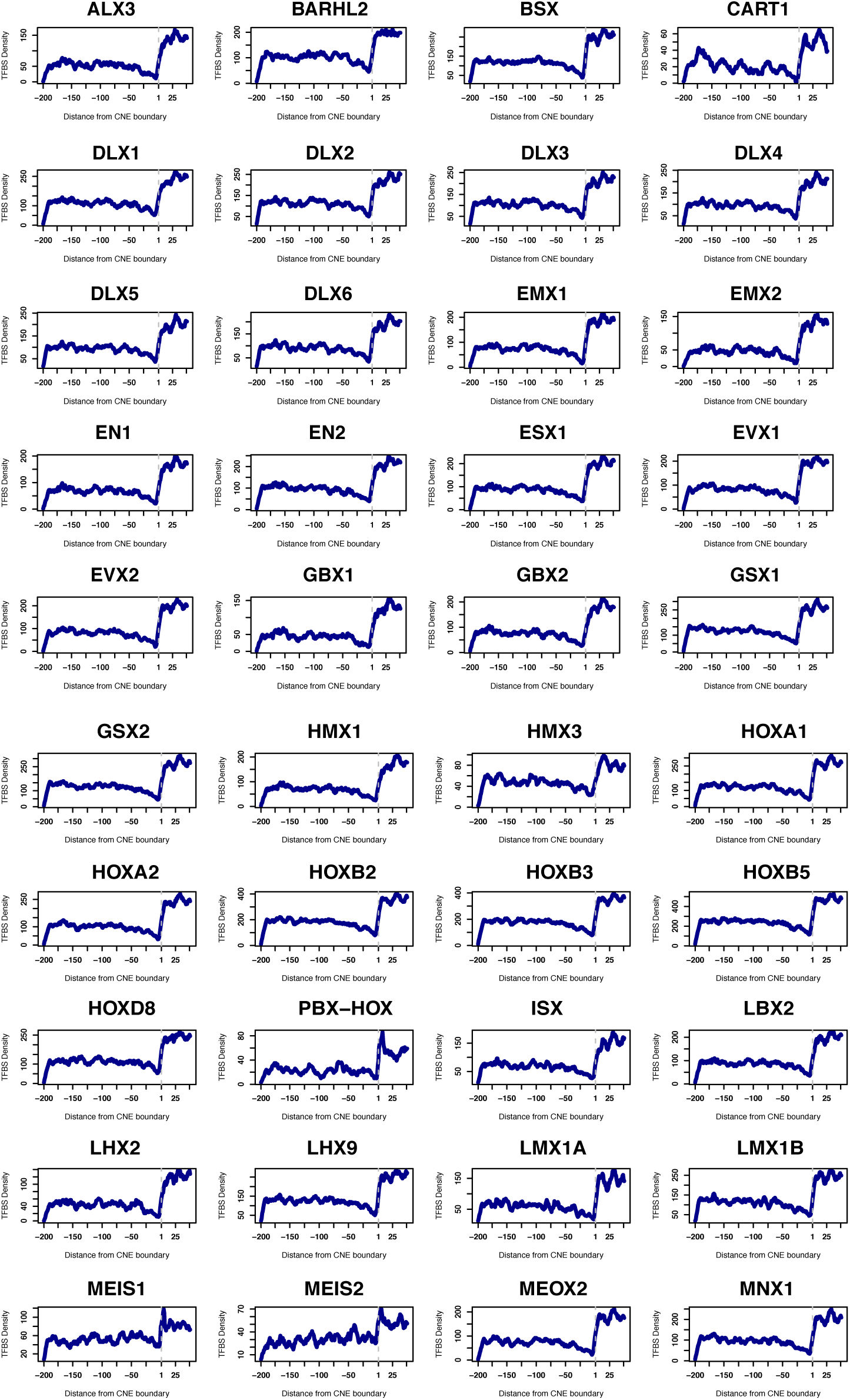
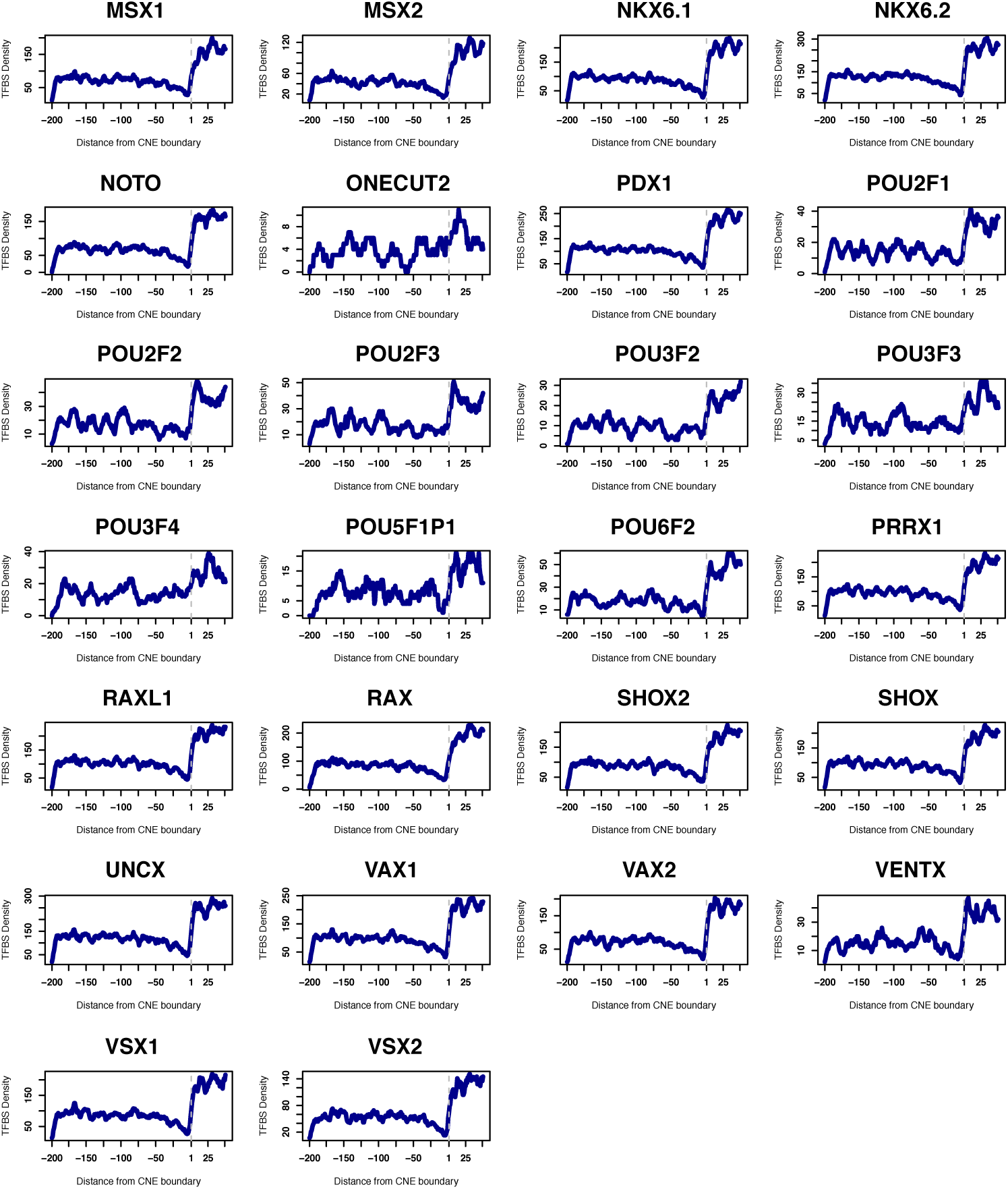
Density of transcription factor binding prediction for 70 individual homeobox containing PWMs, 200bp outside and 50bp inside of CNEs.(Part 2 of 2). All of the homeobox PWMs show increased presence inside of the CNEs, with most showing increasing density as we move towards the center of the CNE. A few PWMs, show very precise enrichment at the very edges of the CNEs including PBX-HOX, MEIS1/2, POU2F1, POU2F2 and POU2F3.

**Table S1:**
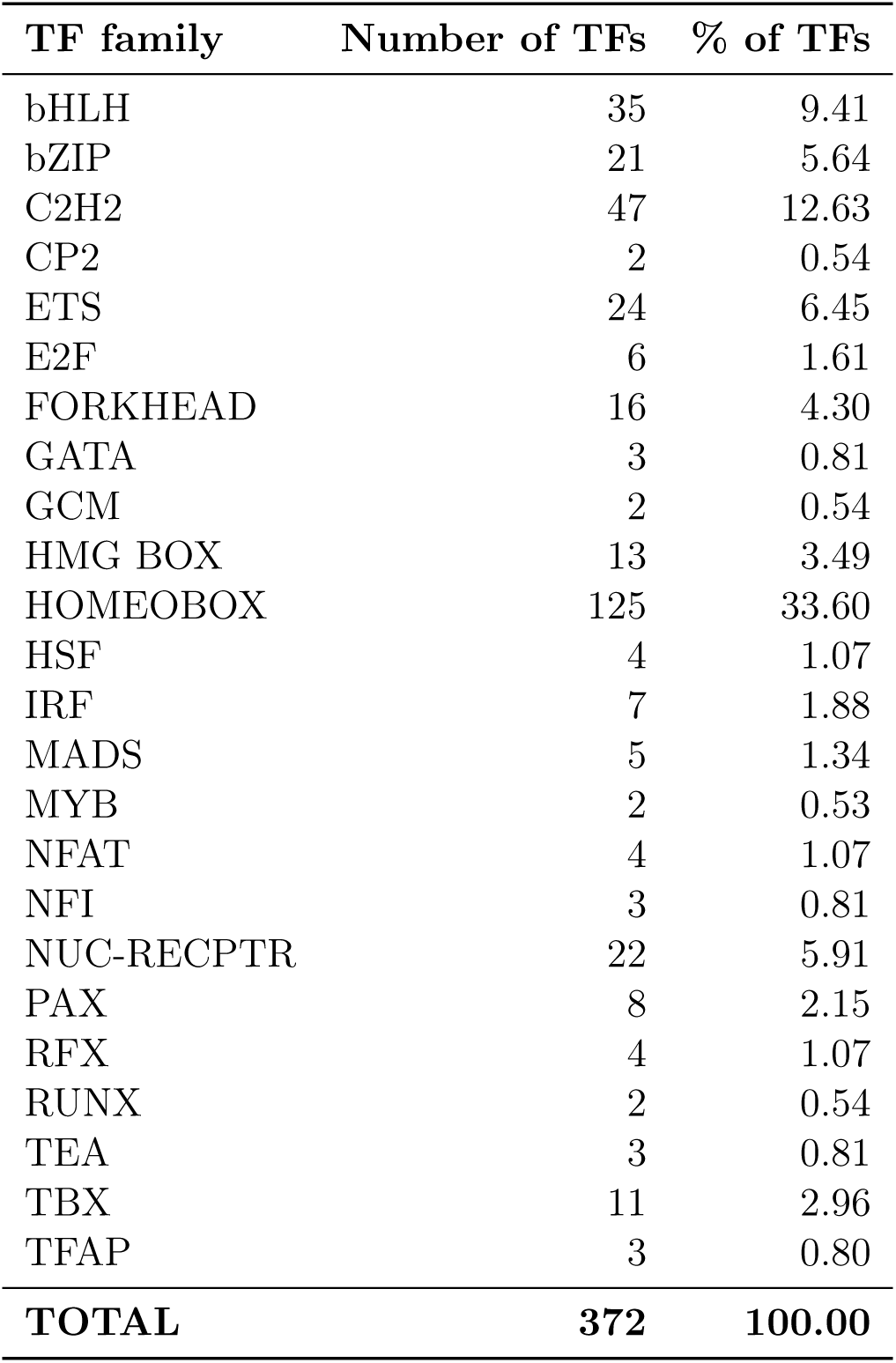
Human Transcription Factor PWMs.

